# An optimized derivative of an endogenous CXCR4 antagonist prevents atopic dermatitis and airway inflammation

**DOI:** 10.1101/2020.08.28.272781

**Authors:** Mirja Harms, Monica MW Habib, Simona Nemska, Antonella Nicolò, Andrea Gilg, Nico Preising, Pandian Sokkar, Sara Carmignani, Martina Raasholm, Gilbert Weidinger, Gönül Kizilsavas, Manfred Wagner, Ludger Ständker, Ashraf Abadi, Hassan Jumaa, Frank Kirchhoff, Nelly Frossard, Elsa Sanchez-Garcia, Jan Münch

## Abstract

**Background:** Aberrant CXCR4/CXCL12 signaling is involved in many pathophysiological processes including chronic inflammatory diseases. Thus, the chemokine receptor CXCR4 is a promising target for the therapy of inflammatory disorders, such as atopic dermatitis or allergic asthma. A natural fragment of serum albumin, named EPI-X4, has previous been identified as endogenous peptide antagonist and inverse agonist of CXCR4. The endogenous CXCR4 antagonist provides a promising basis for the development of improved analogues for the therapy of inflammatory diseases.

**Objective:** To increase the anti-CXCR4 activity of EPI-X4 and to evaluate the therapeutic potential of optimized analogs in mouse models of atopic dermatitis and asthma.

**Methods:** Molecular docking analysis of the interaction of EPI-X4 with CXCR4 was performed to define critical interaction motifs and to rationally design analogs with increased activity. EPI-X4 derivatives were synthesized and CXCR4 binding and antagonizing activity determined in assays for antibody competition, inhibition of CXCR4-mediated HIV-1 infection, CXCL12-dependent Ca^2+^ mobilization, ERK and AKT phosphorylation and cell migration. Toxicity of peptides was evaluated in zebrafish embryos. The therapeutic efficacy of the lead peptide EPI-X4 JM#21 was determined in mouse models of atopic dermatitis and asthma.

**Results:** Docking analysis identified key interaction motifs of EPI-X4/CXCR4. Rational drug design allowed to increase the anti-CXCR4 activity of EPI-X4 and resulted in the generation of the lead analog JM#21, which bound CXCR4 and suppressed CXCR4-tropic HIV-1 infection more efficiently than the clinically approved small molecule CXCR4 antagonist AMD3100. JM#21 did not exert toxic effects in zebrafish embryos and efficiently prevented inflammation of the skin in a mouse model of atopic dermatitis. Moreover, EPI-X4 and its improved derivative suppressed allergen-induced infiltration of eosinophils and other immune cells into the airways of animals in an asthma mouse model.

**Conclusion:** The rationally designed EPI-X4 derivative JM#21 is a potent antagonist of CXCR4 and the first CXCR4 inhibitor with therapeutic efficacy in atopic dermatitis. Further clinical development of this new class of CXCR4 antagonists for the therapy of atopic dermatitis, asthma and other CXCR4-associated diseases is highly warranted.

**Graphical Abstract:** 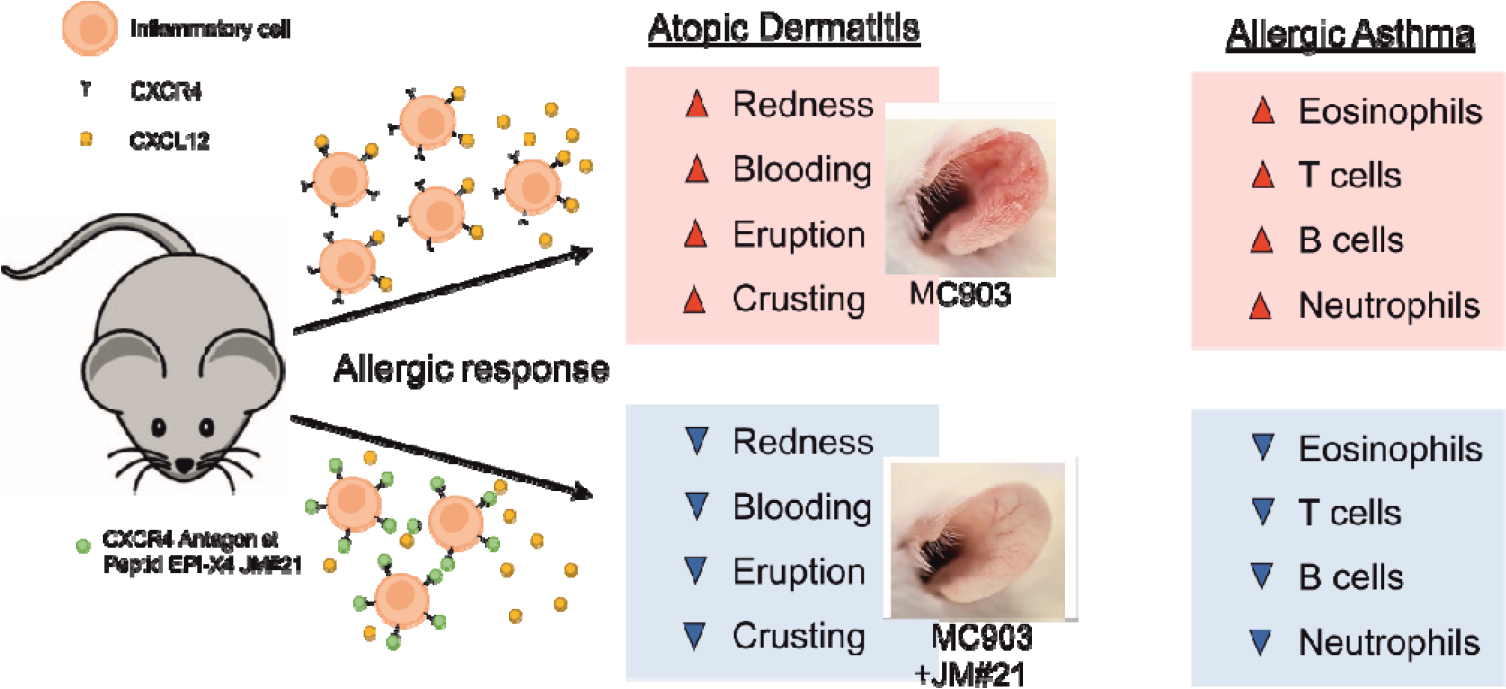

## Introduction

The CXCR4–CXCL12 signaling axis plays a pivotal role in numerous biologic processes, i.e. organ development, hematopoiesis, vascularization, cell and tissue renewal, hematopoietic stem cell retention, inflammation, and immune control^1,2,3,4,5^. In addition, deregulation of this axis e.g. by overexpression of CXCR4, is associated with a variety of diseases, such as tumor development, progression and metastasis, neurodegenerative and autoimmune disorders, as well as immunodeficiencies or inflammatory diseases, including atopic dermatitis and several forms of asthma^6,7,8,9,10,11^. CXCR4 is a long-known drug target and several small molecules, peptide or antibody-based CXCR4 antagonists are currently in preclinical and clinical development^12^. However, the only clinically used CXCR4 antagonist is Plerixafor (Mozobil, AMD3100), which is approved as stem cell mobilizing agent in autologous transplantation of bone marrow cells in patients with Non-Hodgkin’s lymphoma or multiple myeloma, but not suitable for the treatment of chronic disease because of significant adverse effects^13,14^.

One CXCR4/CXCL12-linked disease is asthma, a pathologic overresponse (airway hyper-responsiveness) to a wide variety of stimuli. Inhaled allergens, for example, are detected by pattern recognition receptors on the surface of airway epithelial cells. This triggers secretion of a variety of inflammatory cytokines which stimulate e.g. lung-associated dendritic cells, that migrate into draining lymph nodes and induce a type 2 allergic response by activation of effector cells. Eosinophilia and increased influx of TH2 cells can then be measured in broncho-alveolar lavage (BAL) or bronchial biopsies. Continuous stimuli may lead to chronic inflammation resulting in severe asthma symptoms including increased mucus secretion, thickening of the airway wall and subsequent narrowing of the airway lumen, inducing shortness of breath, wheeze, cough, and chest tightness^15,16,17^. Asthma requires treatment with inhaled corticosteroids plus a second controller. Severe forms are treated with systemic corticosteroids and nowadays also monoclonal antibodies that target proinflammatory cytokines^18^. As in other inflammatory diseases, the CXCR4/CXCL12 axes is involved in asthma pathology: CXCL12 is released in high concentrations into BAL of patients with asthma and correlates with leukocyte infiltration suggesting that it contributes to allergic cell recruitment in asthma^19^. In addition, CXCR4 expression was shown to be upregulated on eosinophils in the BAL of patients with lung eosinophilia^20,21^ and CXCR4 overexpression on leukocytes is associated with stronger inflammatory processes^22,19^. In mouse models of asthma, targeting CXCR4 with an anti-CXCR4 antibody, or AMD3100, or by neutralizing CXCL12 with inhibiting ligands, reduced eosinophilia and migration of immune cells into the airways, and subsequent release of inflammatory cytokines^22–26^.

CXCR4/CXCL12 also plays a role in inflammatory diseases of the skin, in particular psoriasis, as both molecules are upregulated in inflamed areas of the skin in transgenic mice, as well as human biopsies^9,27,28^. Moreover, treatment with AMD3100 and CXCL12 neutralizing antibodies inhibited skin inflammation in mice^27^. However, little is known about the involvement of CXCR4/CXCL12 in atopic dermatitis, a chronically relapsing inflammatory skin disorder with a prevalence of about 1 to 3 % in adults, and 15 to 20 % in children, often accompanied by allergic rhinitis, asthma, food allergy, and conjunctivitis^29^. Inflammation is caused by an array of chemokines and adhesion factors that recruit leukocytes to the skin, where they differentiate and become activated by e.g. dendritic cells, allergens or yet unknown factors^30^. These processes result in disturbed skin barrier function, an elevated and more frequent allergic response, as well as defects in antimicrobial immune defense^31^. Currently, there is no cure for atopic dermatitis. The inflamed skin is treated with topical immunosuppressive corticosteroids and calcineurin inhibitors or, in more severe cases, with oral corticosteroids and dupilumab, a monoclonal antibody that inhibits action of proinflammatory cytokines IL-4 and IL-13^32^. However, prolonged therapy with corticosteroids and calcineurin is associated with side effects and antibody therapy is relatively expensive, necessitating the development of new treatment options^33^.

Collectively, these data show that antagonizing CXCR4 is a seminal approach for the treatment of asthma and chronic inflammatory diseases of the skin. One promising novel CXCR4 antagonist is EPI-X4 (Endogenous Peptide Inhibitor of CXCR4)^34,35,36^. EPI-X4 is a 16-mer peptide derived from human serum albumin (HSA) originally identified as inhibitor of CXCR4-tropic HIV-1^34,37^. It binds to CXCR4 thereby abrogating binding of CXCL12 and the HIV-1 gp120. Interestingly, EPI-X4 not only antagonized CXCL12-induced CXCR4 signaling but also reduced the basal signaling activity of the receptor in the absence of the chemokine, rendering EPI-X4 also an inverse agonist of CXCR4^34^. The peptide is released from albumin by ubiquitous aspartic proteases, but its physiological function is still unclear^37^. Binding of EPI-X4 to CXCR4 is highly specific as the peptide does not interfere with signaling of other chemokine/chemokine receptor pairs^35^. Furthermore, EPI-X4 was shown to block CXCL12-mediated receptor internalization and suppressed the migration and invasion of cancer and immune cells towards a CXCL12 gradient. Studies in mice provided proof-of concept evidence that EPI-X4 has therapeutic potential since the peptide mobilized hematopoietic stem/progenitor cells and inhibited recruitment of inflammatory immune cells into the lung. However, high concentrations of the endogenous peptide were administered (~ 2 mg EPI-X4 per mouse i.p.)^34^. A preliminary structure-activity-relationship (SAR) study allowed to increase the CXCR4 blocking activity of EPI-X4 by about 1.5 to 2 orders of magnitude^34^. We here set out to further optimize the anti-CXCR4 activity of EPI-X4 using rational drug design and to evaluate the therapeutic potential of the lead candidate in mouse models of asthma and atopic dermatitis. We show that the optimized EPI-X4 analog JM#21 binds CXCR4 with high affinity and antagonizes CXCL12-elicited CXCR4 responses *in vitro*. Most notably, topical application of JM#21 prevented inflammation of the skin and lungs in mouse models of dermatitis and asthma.

## Material and Methods

### Molecular modeling

The reported crystal structure of CXCR4 (PDB ID: 3ODU) was used for the modelling studies^38^. The missing N-terminal loop was added using the NMR structure of the N-terminus of CXCR4 (PDB ID: 2K04)^39^. The NMR structure of EPI-X4 (PDB ID: 2N0X)^34^ was used for the docking studies with CXCR4. The molecular docking calculations were carried out using the HADDOCK v2.2 webserver available at http://www.bonvinlab.org/software/haddock2.2/^40^.

### Reagents and peptide synthesis

All chemicals were used as provided by the manufacturers. Amino acids were purchased from Novabiochem (Merck KGaA, Darmstadt, Germany). N, N dimethylformamide (DMF), 20 % (v/v) piperidine in DMF, O-benzotriazole-N, N, N’N’-tetramethyluronium-hexafluoro-phosphate (HBTU) and trifluoroacetic acid (TFA) were purchased from Merck Millipore (Merck KGaA, Darmstadt, Germany). Triisopropylsilane (TIS), diisopropylethylamine (DIEA) were purchased from Sigma Aldrich (Sigma-Aldrich Chemie GmbH Munich, Germany). Acetonitrile was purchased from JT.Baker (Avantor Performance Materials B.V. 7418 AM Deventer Netherlands). The peptides were synthesized automatically on a 0.10 mmol scale using standard Fmoc solid phase peptide synthesis techniques with the microwave synthesizer (Liberty blue; CEM). A preloaded resin was used and provided in the reactor. The Resin was washed with DMF. The Fmoc protecting group was removed with 20 % (v/v) piperidine in DMF and initialized with microwave followed by washing with DMF. Amino acids were added in 0.2 M equivalent to the reactor, then HBTU in a 0.5 M equivalent was dosed to the amino acid solution. After that, 2 M equivalent of DIEA was added to the resin. The coupling reaction was proceeded with microwaves for a few minutes then the resin was washed in DMF. These steps were repeated for all amino acids in the sequence. The last amino acid was deprotected. Once the synthesis was completed, the peptide was cleaved in 95 % (v/v) trifluoroacetic acid (TFA), 2.5 % (v/v) triisopropylsilane (TIS), and 2.5 % (v/v) H_2_O for one hour. The peptide residue was precipitated and washed with cold diethyl ether (DEE) by centrifugation. The peptide precipitate was then allowed to dry under air flow to remove residual ether. The peptide was purified using reversed phase preparative high-performance liquid chromatography (HPLC; Waters) in an acetonitrile/water gradient under acidic conditions on a Phenomenex C18 Luna column (5 mm pore size, 100 Å particle size, 250 _ 21.2 mm). Following purification, the peptide was lyophilized on a freeze dryer (Labconco) for storage prior to use. The purified peptide mass was verified by liquid chromatography mass spectroscopy (LCMS; Waters). AMD3100 octahydrochloride hydrate was purchased from Sigma Aldrich (#A5602) and dissolved in H_2_O. CXCL12 was purchased from Peprotech (#300-28A) and dissolved at a concentration of 100 μg/mL in H_2_O.

### Nuclear Magnetic Resonance Spectroscopy

Nuclear Magnetic Resonance spectra were collected on a Bruker 700 MHz AvanceIII spectrometer with a 5 mm BBI ^1^H/BB probe equipped with a z-gradient and on a 850 MHz AVANCE III system equipped with a 5 mm triple resonance TXI ^1^H/^13^C/^15^N probe with a *z*-gradient at 298 K. NMR samples with JM#21 or WSC02 were prepared with 5 mg peptide each in 500 μl 20 mM sodium phosphate buffer pH 7.4 containing 5 % (v/v) D_2_O. NMR samples with WSC02 were prepared with 5 mg WSC02 in 500 μl MilliQ-water containing 5 % (v/v) D_2_O. Several homo- and heteronuclear 2D NMR spectra (HSQC, HMBC, COSY, TOCSY, and NOESY) were recorded for the assignment of the ^1^H and ^13^C resonances at natural isotope abundance. NMR spectra were processed using Topspin 3.6 and were analyzed using NMRFAM-SPARKY^41^. After manual peak assignment and extraction of the peak intensities, the data was incorporated into the program ARIA for structure calculations^42^.

### Cell culture

TZM-bl HIV-1 reporter cells stably expressing CD4, CXCR4 and CCR5 and harboring the lacZ reporter genes under the control of the HIV LTR promoter were obtained through the NIH AIDS Reagent Program, Division of AIDS, NIAID, NIH: TZM-bl cells (Cat#8129) from Dr. John C. Kappes, and Dr. Xiaoyun Wu. TZM-bl cells and HEK293T cells were cultured in DMEM supplemented with 10 % fetal calf serum (FCS), 100 units/mL penicillin, 100 μg/mL streptomycin, and 2 mM L-glutamine (Gibco). SupT1 cells were cultured in RPMI supplemented with 10 % FCS, 100 units/mL penicillin, 100 μg/mL streptomycin, 2 mM L-glutamine and 1 mM HEPES (Gibco). The BCR-ABL cell line was generated introducing the BCR-ABL expression vector into bone marrow-derived wild type pro/pre B cells. Cells were cultured in Iscove’s basal medium supplemented with 10 % FCS, 100 U/ml Penicillin, 100 U/ml Streptomycin, 2mM L-glutamine and 50 μM β-mercaptoethanol.

### HIV-1 inhibition assay

Viral stocks of CXCR4-tropic NL4-3 were generated by transient transfection of HEK293T cells with proviral DNA as described before^34^. Inhibition of viral infection was performed in TZM-bl reporter cells. For this 70 μL of 1 x 10^5^ cells (in growth medium supplemented with 2.5 % FCS) were pretreated with 10 μL of inhibitors for 15 min at 37°C. Cells were then inoculated with 20 μL of diluted virus. Infection rates were determined after 3 days using Gal-Screen system (Applied Biosystems).

### Ca^2+^ mobilization assay

A well-established murine cell line expressing the fusion protein BCR-ABL was used for the experiments. Measurement of intracellular calcium flux was performed as described previously^43^. 1×10^6^ cells were incubated with 5□μg/mL of Indo□1 AM (Invitrogen) and 0.5□μg/mL of pluronic F□127 (Invitrogen) in Iscove’s medium supplemented with 1 % FCS (Pan Biotech) at 37°C for 4□min. Cells were then pelleted by centrifugation and resuspended in Iscove’s medium with 1 % FCS and treated with the EPI-X4 derivatives for 10 min at 37°C. Prior measurement cells were pre-warmed for 5 min at 37°C. Calcium flux was assessed by FACS measurement at BD LSR Fortessa. After 30 sec of baseline recording, CXCR4-dependent calcium signaling was determined by stimulating with 100 ng/ml of mouse SDF-1α (PeproTech). To quantify CXCL12-induced Ca^2+^ mobilization, the kinetics analysis platform of FlowJo analysis software (BD Biosciences) was used to determine the area under the curve (AUC) of each calcium flux plot. In order to allow for inter-experimental comparison, the AUC was calculated after baseline subtraction for each single plot. AUC values were exported from FlowJo into MS-Excel where calculations were performed. Statistical analysis was performed using GraphPad Prism 8. One-way ANOVA with Tukey‘s multiple comparisons test was performed. Degree of significance is indicated as follows: \**p* < 0.05; \*\**p* < 0.01; \*\*\**p* < 0.001; ns = not significant.

### Erk and Akt signaling

CXCL12 induced phosphorylation of Erk and Akt was monitored by phospho flow cytometry as described before^44^. For this 100.000 SupT1 cells were starved for 2 h at 37°C (growth medium supplemented with 1 % FCS). Inhibitors were then added for 10 min and cells subsequently stimulated with 100 ng/mL CXCL12 for 2 min. The reaction was stopped by adding 1 % PFA and shifting to 4°C for at least 10 min. Cells were then permeabilized with ice cold methanol and stained with phospho-p44/42 MAPK (Erk1) (Tyr204)/ (Erk2) (Tyr187) (D1H6G) mouse mAb (cell signaling, #5726) and phospho-Akt (Ser473) (193H12) rabbit mAb (cell signaling, #4058) and adequate secondary antibodies for flow cytometry. Signal in the unstimulated control without inhibitor was defined as background and set to 0 %. CXCL12 induced signal without inhibitor was set 100 %.

### Cell migration assay

Migration assays towards a 100 ng/mL CXCL12 gradient were performed with SupT1 cells using 96-well transwell assay plates (Corning Incorporated, Kennebunk, ME, USA) with 5 μm polycarbonate filters. First, 50 μl (0.75 x 10^5^) SupT1 cells resuspended in assay buffer (RPMI supplemented with 0.1 % BSA) were seeded into the upper chamber in the presence or absence of compounds and allowed to settle down for around 15 min. In the meantime, 200 μl assay buffer supplemented with or without 100 ng/ml CXCL12 as well as compounds were filled into a 96 well-V plate. Cells were allowed to migrate towards CXCL12 by putting upper chamber onto the 96 well-V plate. After a migration time of 4 h at 37°C (5 % CO_2_) lower compartment were analyzed for cell content by Cell-Titer-Glo^®^ assay (Promega, Madison, WI, USA). Percentages of migrated cells were calculated as described before^45^ and normalized to the CXCL12-only control.

### Antibody competition assay

Competition with the ECL-2 specific CXCR4 antibody was performed as described before^44^. Compounds were serially diluted in cold PBS and afterwards added to 50,000 SupT1 cells. APC-conjugated anti-human CXCR4 antibody (clone 12G5, #555976, BD) was diluted in PBS containing 1 % FCS and added immediately afterwards at a final concentration of 0.245 nM. After 2 h incubation at 4°C, unbound antibody was removed and cells analyzed in flow cytometry.

### Toxicity in zebrafish

Wild-type zebrafish embryos were dechorionated at 24 h post fertilization (hpf) using digestion with 1 mg/ml pronase (Sigma) in E3 medium (83 μM NaCl, 2.8 μM KCl, 5.5 μM CaCl_2_, 5.5 μM MgSO_4_). Embryos were exposed for 24 h in groups of 3 to 100 μl of E3 containing the test substances. Peptides were tested at 3, 30 and 300 μM. Each concentration was tested in two independent assays, each of which was performed on 10x 3 embryos. The peptide solvent (PBS), diluted in E3, was used as negative control at the same amount as introduced by the highest peptide concentration. As positive control for acute toxicity the pleurocidin antimicrobial peptide NRC-03 (GRRKRKWLRRIGKGVKIIGGAALDHL-NH2) was used at a concentration of 6 μM^46^. At 48 hpf (after 24 h of incubation) embryos were scored in a stereomicroscope for signs of acute toxicity (lysis and/or necrosis), developmental toxicity (delay and/or malformations), or cardiotoxicity (heart edema and/or reduced or absent circulation). Chi-Square test was used to calculate whether the distribution of embryos into toxicity classes differed significantly between the PBS negative control and the test substances.

### Mice

Nine-week-old male Balb/c mice (Janvier, France) were used. The animals were maintained on a 12/12-hour light/dark cycle, with food and water *ad libitum*. All animals received humane care in compliance with the guidelines formulated by the French Ministry of Agriculture and of Higher Education and Research, who authorized the procedures (APAFiS #10067 and #1341).

### Effect of EPI-X4 JM#21 in a mouse model of atopic dermatitis

Mice were anesthetized by isoflurane 5 % (Dechra) before each topical application. MC903 (calcipotriol hydrate, Sigma) was dissolved in EtOH 100 % and topically applied at 2.5 nmol/ear every two days from day (D) D0 to D12 on both sides of the mouse ears. EtOH 100 % was applied to control mice every two days. All peptides were solubilized in EtOH 80 % and applied topically at 400 nmol/ear (= 16μmol/kg/ear) in a volume of 25 μL/ear every day from D0 to D12, 1h after MC903 application. Reference compound dexamethasone (Dex, Sigma) was also used topically and daily at 20 nmol/ear (= 0.8 μmol/kg/ear) and dissolved in EtOH 80 %. The reference CXCR4 non peptidic antagonist AMD3100 (Euromedex) was used daily, topically at 315 nmol/ear (= 12.6 μmol/kg/ear) and dissolved in EtOH 80 %. Ear thickness was measured with a digital micrometer every day before the next topical application, and a mean value of the measurements of both ears was calculated. Scores were registered daily to evaluate the severity of the atopic dermatitis, following the scoring system on Table 1. Photographs of the ears were taken at D12.

**Table 1:** Scoring system for atopic dermatitis model

| Clinical signs | redness | bleeding | eruption | crusting | SUM |
| --- | --- | --- | --- | --- | --- |
| *Severity* |  |  |  |  |  |
| *absent* | 0 | 0 | 0 | 0 | **0 / score min** |
| *mild* | 1 | 1 | 1 | 1 | **4** |
| *moderate* | 2 | 2 | 2 | 2 | **8** |
| *severe* | 3 | 3 | 3 | 3 | **12 / score max** |

### Effect of EPI-X4 JM#21 in a mouse model of allergic airway hypereosinophilia

Mice were sensitized by intraperitoneal (i.p.) injection of a mixture containing 50 μg ovalbumin (OVA, grade V, Sigma) and 2 mg Al(OH)_3_ (alum, Sigma) in saline on D0, D1 and D2. On D5, D6 and D7, mice were treated with peptides (16 μmol/kg), AMD3100 (12.6 μmol/kg, Euromedex) or solvent (NaCl 0.9 %) by intranasal administration (i.n.) (12.5 μL/nostril). Two hours later, mice were challenged i.n. with 10 μg of OVA in saline. The unchallenged controls were administered i.n. with saline. Before each intranasal administration, mice were anesthetized with ketamine (75 mg/kg, Merial) and xylazine (5 mg/kg, Bayer). Mice were kept on a hot plate (37°C) until complete awakening. On D8, mice were anesthetized with ketamine (300 mg/kg, Merial) and xylazine (20 mg/kg, Bayer) and bronchoalveolar lavage performed with PBS-EDTA 2 mM^24^. Briefly, a catheter was inserted into the trachea, and lungs were washed 10 times with 500 μL of PBS-EDTA 2 mM. BAL fluid was centrifuged (300 x *g* for 5 min, 4°C), cell pellet suspended in 500 μL of PBS-EDTA and total cells counted on a Muse^®^ Cell Analyser (Millipore) using anti-CD45-PE antibody (eBioscience, 1:500). Differential cell counts were assessed by flow cytometry (LSRII^®^ cytometer, BD Bioscience). BAL cells were added with Fc Block (BD Bioscience) for 20 min and then marker antibodies were incubated for 30 min: CD11c-FITC (BD Bioscience, 1:250), GR-1-PE-eFluor610 (eBioscience, 1:500), CD11b-APC-Cy7 (BD Bioscience, 1:200), CD45-AlexaFluor700 BioLegend, 1:200), CD3-BV650 (BD Bioscience, 1:250), CD19-BV605 (BD Bioscience, 1:250), DAPI (BD Bioscience, 1:500). Live leukocytes were identified as CD45^+^DAPI^-^ cells then differentiated into T cells (CD11b^low^CD19^-^CD3^+^), B cells (CD11b^low^CD19^+^CD3^-^), eosinophils (CD11b^+^CD11c^-^GR1^-^), neutrophils (CD11b^+^CD11c^-^ GR1^+^) and macrophages (CD11b^+^GR1^-^CD11c^+^).

### Calculations and statistical analysis

Statistical analysis was performed in GraphPad Prism (version 8.3.0). IC_50_ curves were determined by nonlinear regression. Differences between groups were tested for statistical significance using a one-way ANOVA followed by Bonferroni’s post-test. Data were considered significantly different when *p* ≤ 0.05.

## Results

### Molecular docking of EPI-X4 binding to CXCR4

Our previous SAR analysis allowed to increase the CXCR4 antagonizing activity of EPI-X4 (IC_50_ in the HIV-1 inhibition assay of ~ 5.9 μM) by approximately 23-fold, resulting in the first-generation lead WSC02 with an IC_50_ of 254 nM (Table 2,^34^). Compared to the parental peptide, the WSC02 analog is C-terminally truncated by four residues and contains four amino acid substitutions (L1I, Y4W, T5S and Q10C). The Cysteine at position 10 was introduced to enable dimerization or coupling reactions to linker, fatty acid or other moieties. To further improve the CXCR4 antagonizing activity, we first aimed at obtaining an overview of the interactions of EPI-X4 with the receptor at the molecular level. For this purpose, we modeled the structure of EPI-X4 bound to CXCR4 using protein-peptide docking. The resulting protein-peptide complex shows that EPI-X4 binds in the extracellular pocket of CXCR4 (Fig. 1a). EPI-X4 establishes salt-bridge interactions with negatively charged residues of CXCR4 via the positively charged amino acids L1, K6 and K7. Thus, the positively charged N-terminus (L1) of EPI-X4 interacts with E31 of CXCR4 (Fig. 1b). K6 and K7 form salt-bridges with D187 and D262 of CXCR4, respectively, and the nonpolar amino acid V2 of EPI-X4 establishes hydrophobic contacts with F29 and A180 of CXCR4.

**Figure 1:**
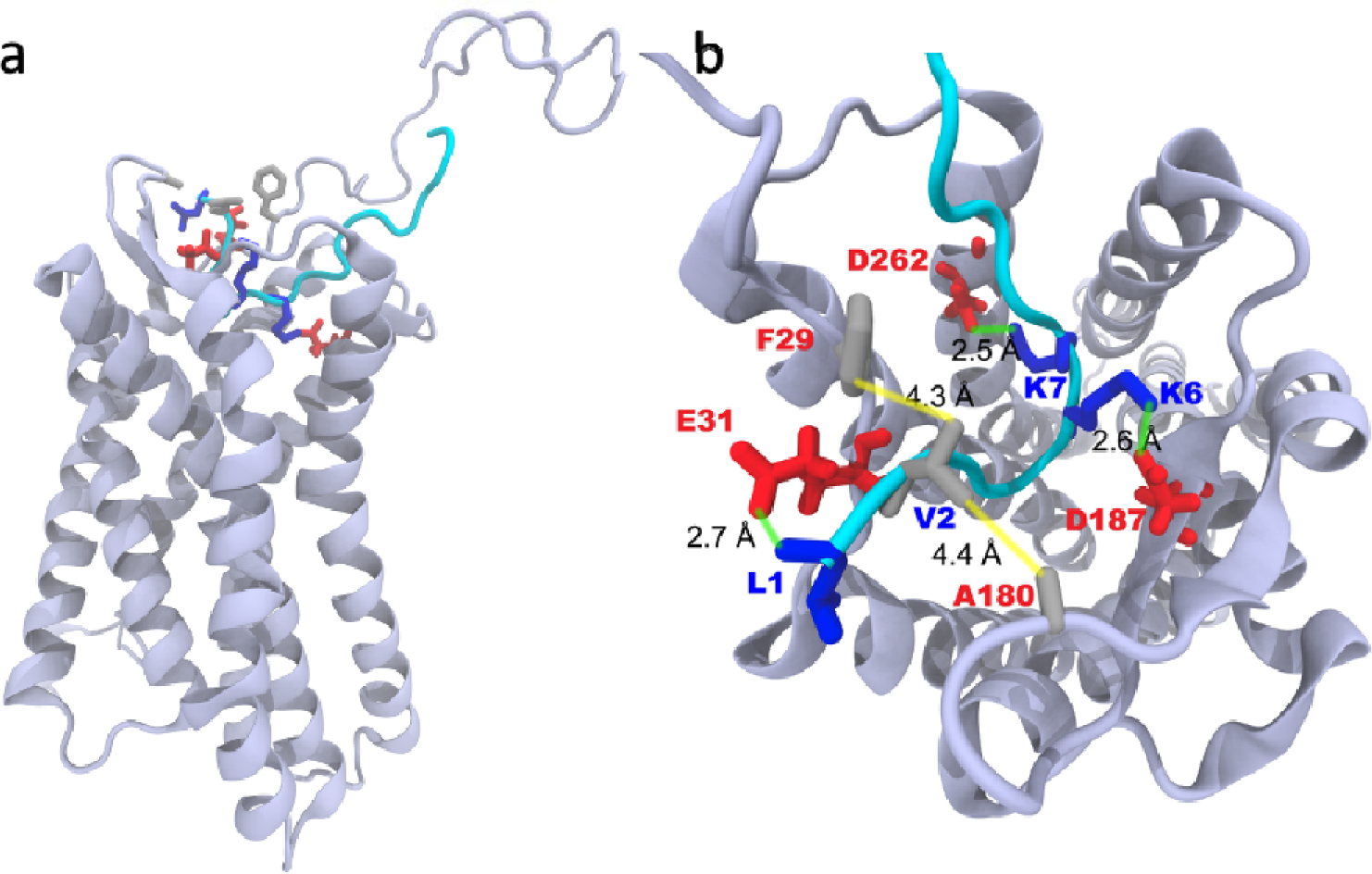
Binding mode of EPI-X4 with CXCR4 obtained by molecular docking. a) Position of EPI-X4 (cyan) in the binding pocket of CXCR4 (cartoon). b) Important interactions that stabilize the protein-peptide complex: L1 (N-terminal amine), K6 and K7 of EPI-X4 are involved in salt-bridge interactions (indicated by green lines) with E31, D187 and D262, respectively. Hydrophobic interactions (indicated by yellow lines) are established between V2 of EPI-X4 and F29 and A180 of CXCR4.

**Table 2.**
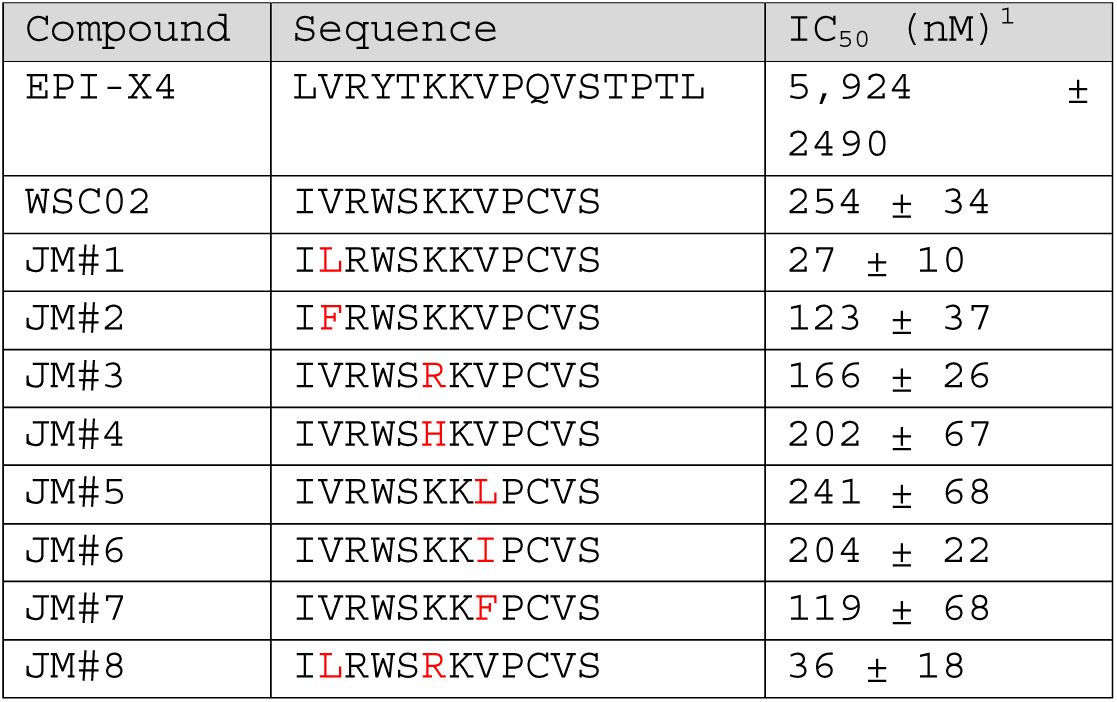
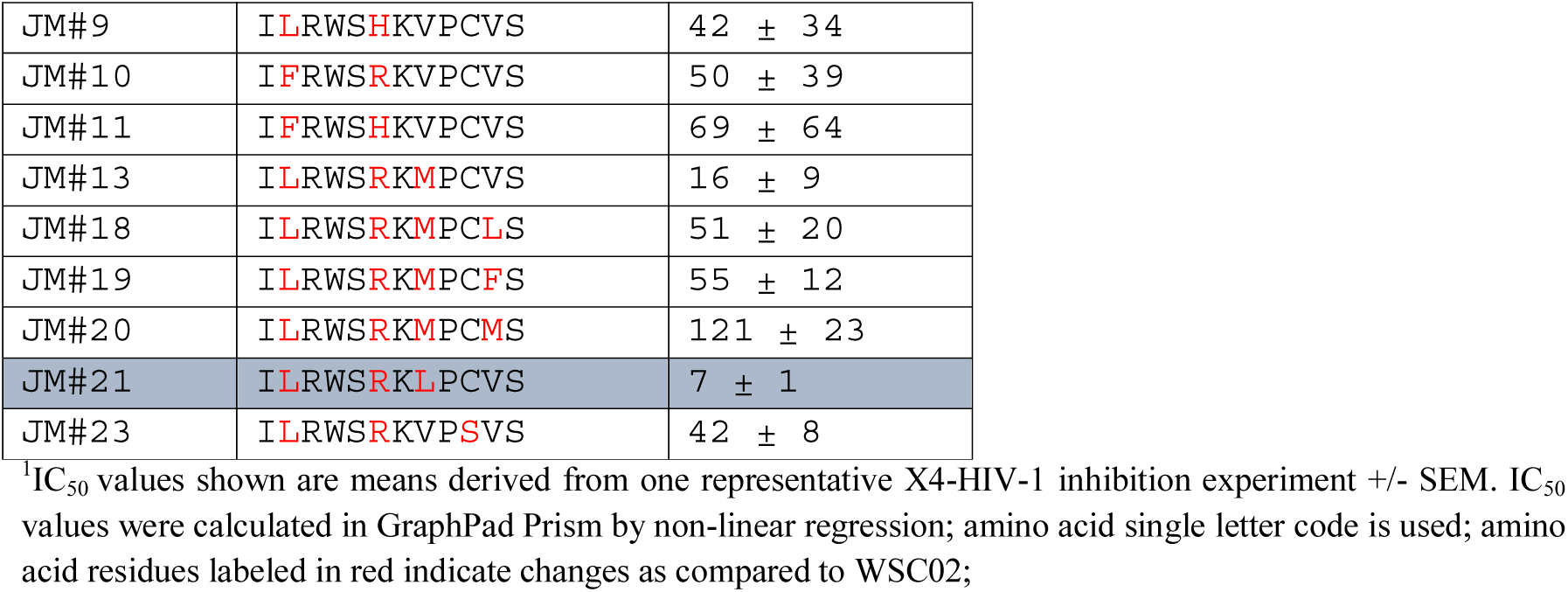
Structure activity relationship study using WSC02 as a template

### Generation of EPI-X4 WSC02 analogs with increased anti-CXCR4 activity

The docking model suggests that mutation at position 2 to a slightly longer aliphatic chain (such as V2>L) or to a relatively large non-polar sidechain (such as V2>F) might improve the hydrophobic contacts between the peptide and the protein. As mentioned above, K6 was also found to be involved in salt-bridge interactions with CXCR4. This supports the hypothesis that changing this amino acid by arginine would increase the binding affinity, since the guanidinium group of arginine can interact strongly with carboxylate groups (via bidentate H-bonds), as compared to the ammonium group. Thus, these and other amino acid substitutions at position 2 and 6 were introduced in WSC02 in order to enhance anti-CXCR4 activity (Fig. 2 a, Table 2). The CXCR4 antagonizing activity of this new set of EPI-X4 peptides (JM#1 – JM#7) was then determined in a CXCR4-tropic HIV-1 infection assay, as described^34^. We used this assay as primary screening test because i) it allows the accurate and fast determination of IC_50_ values, and ii) the anti-HIV-1 activity correlates well with the activity in T-cell migration assays, which are more elaborate^34^. We found, that replacement of the aliphatic and hydrophobic valine (V) at position 2 by leucine (L), harboring a longer functional side-chain (JM#1), increased the antiviral activity of WSC02 by 9-fold (IC_50_ = 27 nM). Substitution by the aromatic phenylalanine (F) had a smaller beneficial effect (JM#2) (IC_50_ = 123 nM). Also, the replacement of the positively charged lysine (K) at position 6 by the positively charged arginine (R) increased the activity of WSC02 (JM#3) (IC_50_ = 166 nM). Replacement of K6 with the weakly basic amino acid histidine (H), however, had no enhancing effect (JM#4) (IC_50_ = 202 nM). Substitution of V at position 8 by either L (JM#5) or isoleucine (I) (JM#6) did not significantly alter antiviral activity (IC_50_ values of 241 nM and 204 nM, respectively). However, substitution with F increased the anti-HIV-1 activity about 2-fold (JM#7) (IC_50_ = 119 nM) (Fig. 2a, Table 2).

**Figure 2.**
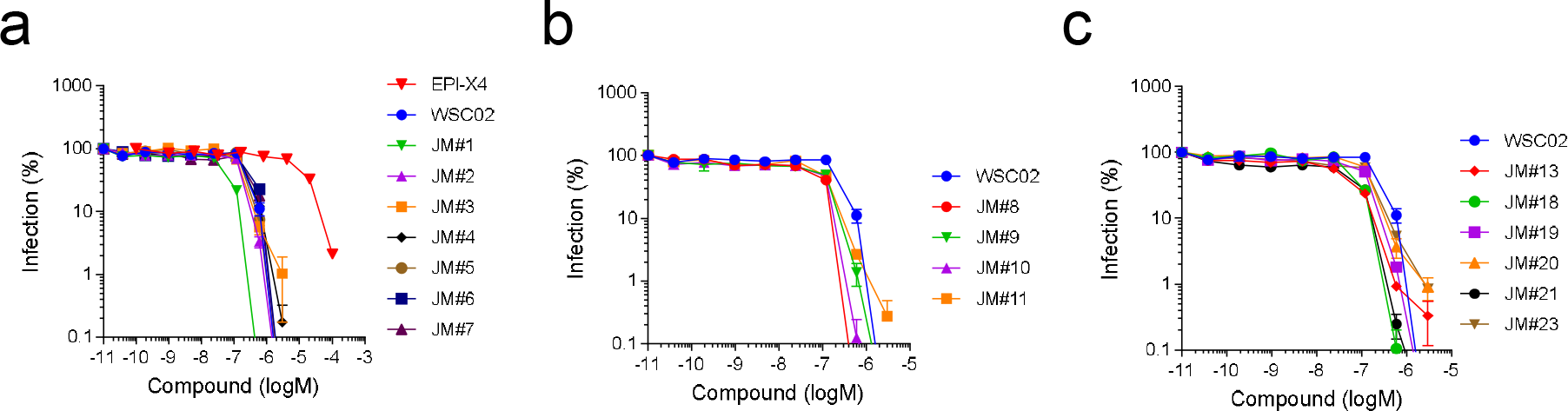
EPI-X4 WSC02 analogues inhibit X4-tropic HIV-1 infection with increased potency. WSC02 derivatives with either a single amino acid substitution (a), two distinct amino acid substitutions (b) or a combination of amino acid substitutions (c) were tested against X4-tropic HIV-1. TZM-bl reporter cells were preincubated with serially diluted peptide and afterwards inoculated with the virus. Infection rates were determined by β-galactosidase assay 3 days later. Shown are data derived from one representative experiment performed in triplicates ± SEM.

Combining beneficial substitutions (JM#8 to JM#11) resulted in analogs with antiviral activities in the two-digit nanomolar range, with JM#8 (V2L and K6R) as the most active analog (IC_50_ = 36 nM) (Fig. 2b, Table 2). Subsequently, based on those findings, additional amino acid substitutions were introduced (JM#13 – JM#23) (Fig. 2c). First, the V at position 8 in JM#8 was replaced by methionine (M) (JM#13) or L (JM#21). Those two derivatives were even more active with IC_50_ values of 16 nM and 7 nM, respectively Based on the sequence of JM#13, V at position 11 was further replaced by either L, F, or M (JM#18 – JM#20) with no beneficial effects. Finally, cysteine (C) 10 in JM#8 was substituted with non-reactive serine. The resulting JM#23 analog was about as active as JM#8 (42 vs 36 nM) suggesting that C10 is not involved in receptor binding (Fig. 2c, Table 2). EPI-X4 analog JM#21 consistently blocked HIV-1 infection with the lowest IC_50_ of 7 ± 1 nM. Thus, it was selected as lead candidate for further characterization.

### 3D NMR-derived solution structure of JM#21 and WSC02

We next analyzed the secondary structures of WSC02 and JM#21 by NMR. The ten best fitting calculated structures of WSC02 (Fig. 3a) and JM#21 (Fig. 3b) based on NMR experimental data, have a backbone root-mean-square deviation (rmsd) of 1.16 +/- 0.33 and 0.83 +/- 0.21, respectively. WSC02 (Fig. 3c) shows a loop formed by the complete peptide-chain. This loop involves, in the best fitting state, a hydrogen bond interaction between the sidechains of S12 and W4 (Fig. 3c). Additionally, the carboxylate group of S12 can form a salt-bridge with the amino group of I1. In contrast, JM#21 shows a C-terminal loop, which is, again in the best fitting structural state, based on an interaction between the backbone carboxylate of S12 and the sidechains of R6, K7, L8 and R3, leaving the N-terminus free (Fig. 3e). The fragment W4-S5-R6 exhibits a turn which can be classified as an α-turn with four peptide-bonds being involved (Fig. 3d). The superimposition of the two peptides shows that WSC02 and JM#21 are slightly similar to each other (backbone rmsd = 2.78 Å) (Fig. 3e), with the main difference that the optimized EPI-X4 JM#21 contains a flexible N-terminus which might facilitate receptor binding to the extracellular pocket (Fig. 1).

**Figure 3:**
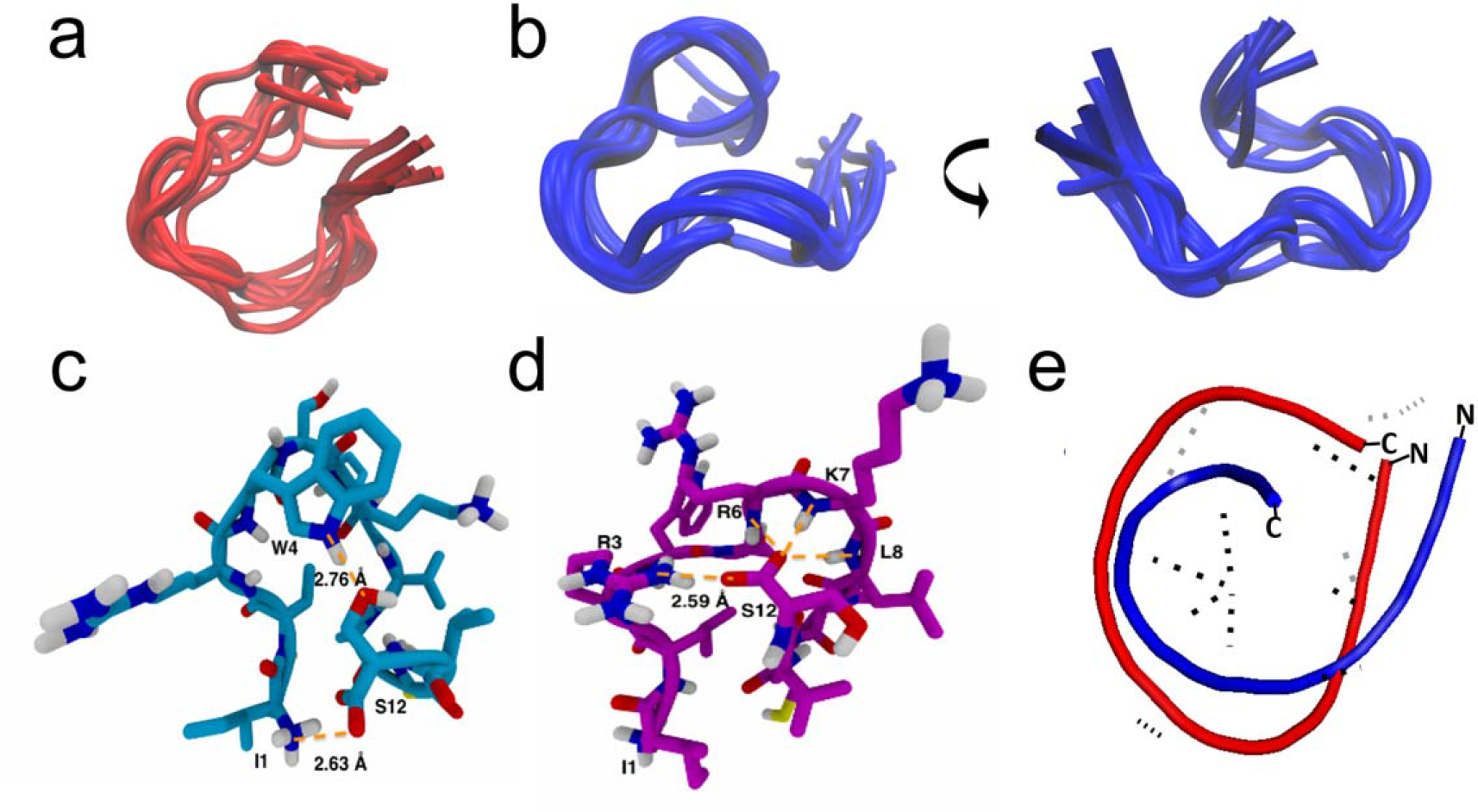
Solution structure of JM#21 and WSC02. Superimposition of the WSC02 (a) and JM#21 (b) conformers with the lowest final target function values calculated with ARIA. The visualization was performed with the Molecular Graphics Viewer of Visual Molecular Dynamics (VMD)^47^. c + d) Intramolecular H-bonds in the WSC02 (c) and JM#21 (d) peptides. (Color code: C = pink/blue, H = white, N = Blue, and O = red. H-bonds are shown as dotted lines in orange). e) Superimposed structures of WSC02 (red) and JM#21 (blue).

### EPI-X4 JM#21 is a potent CXCR4 antagonist

We next determined binding affinities of EPI-X4 and analogs to CXCR4 using a recently described CXCR4 antibody competition assay^44^. This assay is based on the observation that most if not all CXCR4 antagonists interact with CXCR4 in a way that binding of the CXCR4 12G5 antibody raised against the extracellular loop 2 of the receptor is prevented. As shown in Figure 4, the first (WSC02) and second-generation (JM#21) EPI-X4 analogs competed with 12G5 binding even more efficiently than AMD3100. JM#21 (IC_50_ = 223 nM) was ~ 2-fold more active than WSC02 (IC_50_ = 424 nM) and about 3-fold more active than AMD3100 (IC_50_ = 687 nM) (Table 3). For EPI-X4 an IC_50_ value of 3.24 μM was determined (Table 3).

**Figure 4.**
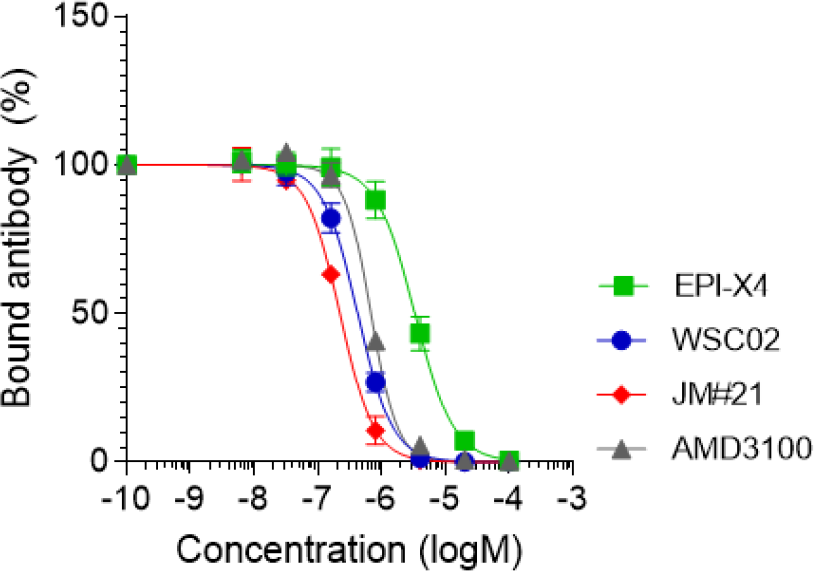
Competition of EPI-X4, WSC02, JM#21 and AMD3100 with the CXCR4 ECL-2 antibody 12G5. CXCR4 expressing SupT1 cells were precooled and assay buffer was removed. EPI-X4 and the optimized derivatives WSC02 and JM#21, and the small molecule AMD3100 were diluted in cold PBS and added to the cells. Immediately afterwards, a constant concentration of the APC-labelled CXCR4 antibody (clone 12G5) was added. After 2 h incubation at 4°C unbound antibody was removed and cells analyzed in flow cytometry. Shown are three individual experiments ± SEM. IC_50_ values were calculated by non-linear regression.

**Table 3.** Comparison between JM#21, WSC02, EPI-X4 and AMD3100

| Compound | IC<sub>50</sub> ± SEM (μM) | | | | |
| --- | --- | --- | --- | --- | --- |
|  | 12G5<sup>1</sup> | X4-HIV-1<sup>2</sup> | pERK<sup>3</sup> | pAKT<sup>3</sup> | Migration<sup>4</sup> |
| EPI-X4 | 3.24 ± 0.62 | 5.924 ± 2.49 | > 100 | 67.60 ± 30.63 | 681.63 ± 305.54 |
| WSC02 | 0.42 ± 0.07 | 0.25 ± 0.03 | 15.25 ± 3.97 | 37.97 ± 26.94 | 15.83 ± 3.50 |
| JM#21 | 0.22 ± 0.02 | 0.007 ± 0.001 | 1.54 ± 0.51 | 2.62 ± 1.40 | 0.48 ± 0.08 |
| AMD3100 | 0.69 ± 0.05 | 0.012 ± 0.002 | 3.28 ± 0.45 | 35.02 ± 27.22 | 0.06 ± 0.01 |

<sup>1</sup> Competition with the CXCR4 specific 12G5 antibody. IC<sub>50</sub> values are means from three individual experiments ± SEM. <sup>2</sup> Inhibition of CXCR4-tropic HIV-1 on TZM-bl cells. IC<sub>50</sub> values are means from one individual experiment performed in triplicates ± SEM. <sup>3</sup> Inhibition of CXCL12 induced ERK or AKT phosphorylation. IC<sub>50</sub> values are means from three individual experiments performed in triplicates ± SEM. <sup>4</sup> Inhibition of SupT1 migration towards a CXCL12 gradient. IC<sub>50</sub> values are means from 4 individual experiments performed in triplicates ± SEM.

Calcium ions (Ca^2+^) are important intracellular messengers since they exert regulatory effects on cytosolic enzymes and proteins and are often induced after CXCR4 activation^48^. Thus, in the next set of experiments, the effect of EPI-X4, its analogs and AMD3100 on CXCL12-induced intracellular Ca^2+^ mobilization was analyzed in BCR-ABL transformed bone marrow cells via flow cytometry^49^. As shown in Fig. 5a,b, CXCL12 treatment induced a transient Ca^2+^ response which was entirely disrupted by 100 μM of the JM#21 analog. At the same concentration, WSC02 only suppressed Ca^2+^ release by ~ 80 %, and EPI-X4 was almost inactive at the tested concentrations (Fig. 5a,b). Here, AMD3100 was more active and reduced Ca^2+^ mobilization already at a concentration of 1 μM, and completely abrogated it at a concentration of 10 μM.

**Figure 5.**
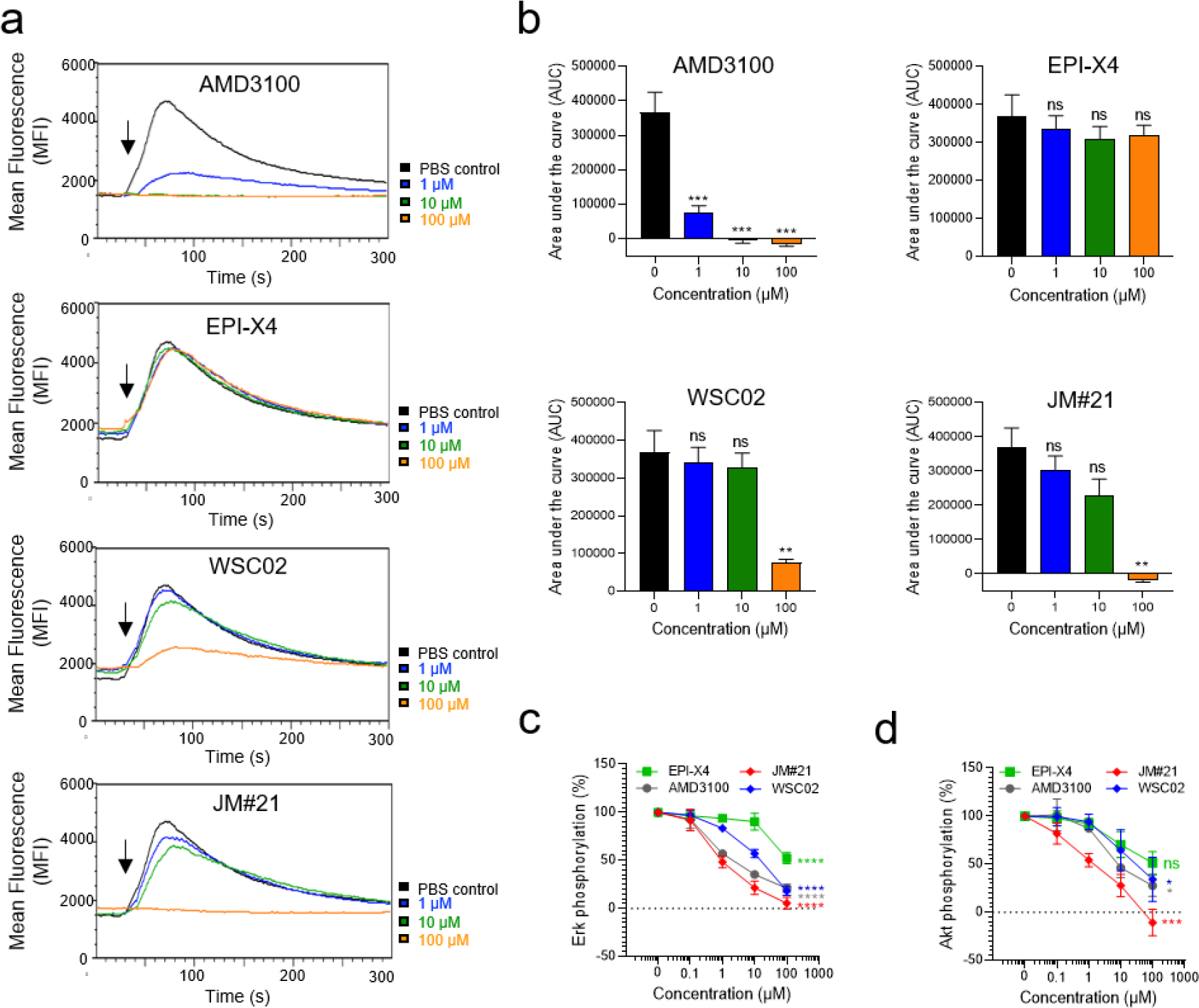
Effect of EPI-X4 derivatives and AMD3100 on CXCL12-evoked Ca^2+^ mobilization and Erk and Akt signaling. (a) Inhibition of CXCL12-induced calcium release. BCR-ABL expressing murine pro/pre B cells were loaded with Indo-1 AM and incubated with inhibitors for 10 min. Baseline fluorescence signal was recorded for 30 sec and calcium flux induced by stimulation with 100 ng/ml mCXCL12 (black arrow). Signal was recorded for additional 260 sec. (b) Areas under the curves (AUC) was calculated after baseline subtraction. Shown are data derived from 3 individual experiments ± SEM. (c, d) Inhibition of CXCL12-induced Erk and Akt phosphorylation. SupT1 cells were preincubated with indicated concentrations of compounds for 15 min in starvation medium and afterwards stimulated with 100 ng/mL CXCL12 for 2 min. Cells were then permeabilized and stained for phosphorylated Erk (c) or phosphorylated Akt (d) and analyzed in flow cytometry. Shown are data derived from 4 individual experiments performed in triplicates ± SEM. IC_50_ values were determined by non-linear regression. \**p*< 0.05; \*\**p* < 0.01; \*\*\**p* < 0.001; \*\*\*\**p* > 0.0001; ns = not significant (one-way ANOVA in comparison to PBS control).

CXCL12 binding to CXCR4 also activates the protein kinase B (PKB, AKT) and extracellular-regulated kinase (ERK) signaling pathways, which are both aberrantly activated in a wide range of human cancers and inflammatory diseases^50–52^. We determined the effect of the CXCR4 antagonists on CXCL12-induced phosphorylation of ERK and AKT in T lymphoblasts by phospho-flow cytometry. JM#21 potently suppressed CXCL12-induced ERK and AKT phosphorylation with IC_50_ values of 1.5 μM (Fig. 5c) and 2.6 μM (Fig. 5d), respectively (Table 3). AMD3100 was slightly less active compared to JM#21 for inhibition of ERK phosphorylation (Fig. 5c) and markedly less potent for AKT phosphorylation with IC_50_ values of 3.3 and 35.0 μM, respectively (Fig. 5d and Table 3). EPI-X4 was least active for inhibition of pERK and pAKT (Table 3).

Finally, the four compounds were analyzed for their ability to block migration of the T lymphoblastic lymphoma cell line SupT1 along a CXCL12 gradient. JM#21 efficiently blocked CXCL12-induced cell migration with IC_50_ values of 0.48 μM (Fig. 6), which was 33-fold more efficient as compared to WSC02 (IC_50_ = 15.8 μM) and more than 1,400-fold more efficient than the wild type peptide EPI-X4 (IC_50_ = 681.6 μM). In this assay, AMD3100 was most potent with an IC_50_ of 0.06 μM (Fig. 5) (Table 3).

**Figure 6.**
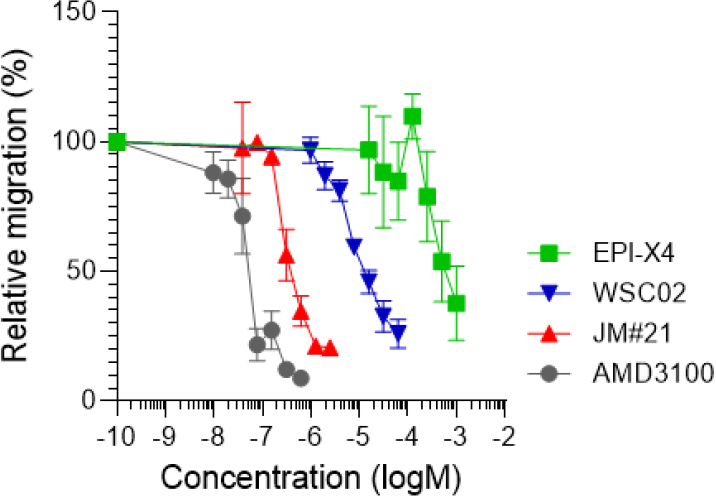
Migration of SupT1 cells in the presence of CXCR4 ligands towards a CXCL12 gradient. The assay was performed in a transwell plate with 5 μm pore size with 100 ng/mL CXCL12 in the lower chamber. The number of migrated cells was determined by CellTiterGlo^®^ assay. Shown are three individual experiments performed in triplicates ± SEM.

### EPI-X4 JM#21 shows no toxicity in zebrafish

Zebrafish embryos provide a useful *in vivo* system for toxicity assessments of chemicals, since their organs and tissues are similar to those of mammals on the molecular, physiological, and basic anatomical level^53^. To evaluate toxicity, zebrafish embryos were exposed for 24 h to a maximum of 300 μM of the peptides or controls, and then analyzed in a stereomicroscope for acute toxicity (lethality, necrosis, lysis), developmental toxicity (developmental delay, malformations) and cardiotoxicity (heart edema, reduced or absent circulation). None of the EPI-X4 analogs induced signs of toxicity whereas the membrane damaging antimicrobial peptide pleurodicin NRC03^46^ was acutely toxic (Fig. 7).

**Figure 7.**
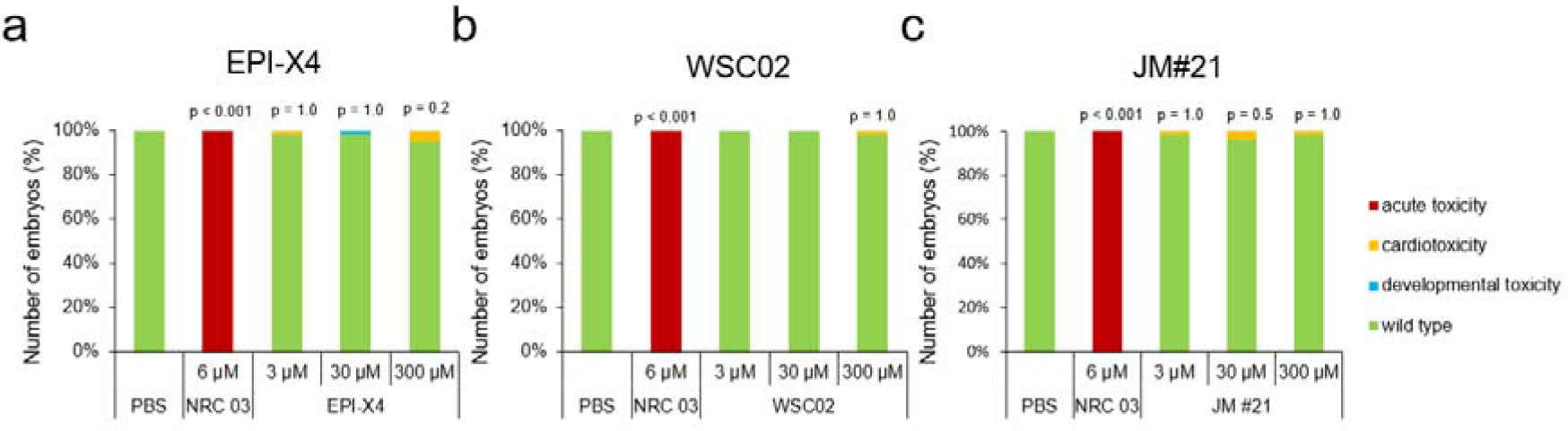
JM#21 is not toxic in an embryonic zebrafish model. At 24 hours post fertilization, dechorionated zebrafish embryos were exposed to EPI-X4 (a), WSC02 (b) or JM#21 (c) for 24 h. As negative control PBS was used, and as positive control the antimicrobial peptide NRC-03. Shown are data derived from 60 embryos per group.

### EPI-X4 JM#21 suppresses airway inflammation in a mouse model of allergic airway hypereosinophilia

It has previously been shown that high doses of EPI-X4 reduce inflammatory cell infiltration in a mouse model of allergic airway hypereosinophilia^34^. To examine whether the EPI-X4 derivative JM#21 may be more potent in preventing airway inflammation, parental EPI-X4 and analogues analogs (16 μmol/kg body weight) and AMD3100 (12.6 μmol/kg body weight) were administered intranasally prior to allergen (OVA) challenge. As controls, mice were not challenged with OVA, and solvent only was administered. At day 8, bronchoalveolar lavage was collected and the number of infiltrated cells determined by flow cytometry (Fig. 8a). Allergen challenge resulted in significant infiltration of eosinophils (Fig. 8c), T cells (Fig. 8d), B cells (Fig. 8e) and neutrophils (Fig. 8f), also demonstrated by the total increase in cell count (Fig. 8b), compared to unchallenged control mice. The parental peptide EPI-X4 inhibited the total cell count by 55 ± 7 % (p < 0.01), the recruitment of eosinophils by 76 ± 7 % (p < 0.01), neutrophils by 63 ± 7 % (p < 0.01), T cells by 61 ± 8 % (p < 0.05) and B cells by 54 ± 9 % (p < 0.05) (Fig. 8b-e). The N-terminally truncated version of EPI-X4 (ALB409-423), which does not antagonize CXCR4^33^, did not interfere with most immune cells, but showed a significant activity in blocking neutrophil infiltration, for unknown reasons. Interestingly, the lead candidate JM#21 was most effective in suppressing migration of immune cells into the airways. Specifically, JM#21 inhibited the recruitment of eosinophils by 80 ± 1 % (p < 0.01), T cells by 73 ± 13 % (p < 0.01), B cells by 71 ± 5 % (p < 0.05), and neutrophils by 42 ± 15 % (p < 0.05), as well as the total inflammatory cell influx by 61 ± 12 % (p < 0.01) (Fig. 8b-f). The control CXCR4 antagonist, AMD3100, also inhibited recruitment of total cells (41 ± 13 %, p < 0.05), eosinophils (56 ± 13 %, p < 0.05), T cells (47 ± 15 %, p < 0.05), B cells (47 ± 17 %, p < 0.05) and neutrophils (57 ± 14 %, p < 0.05), but was on average less active than JM#21. None of the compounds analysed resulted in infiltration of cells in unchallenged mice. Thus, EPI-X4 JM#21 is an effective inhibitor of lung inflammation and particularly potent in preventing infiltration of eosinophils, which are believed to play a key role in asthma exacerbations^54^.

**Figure 9.**
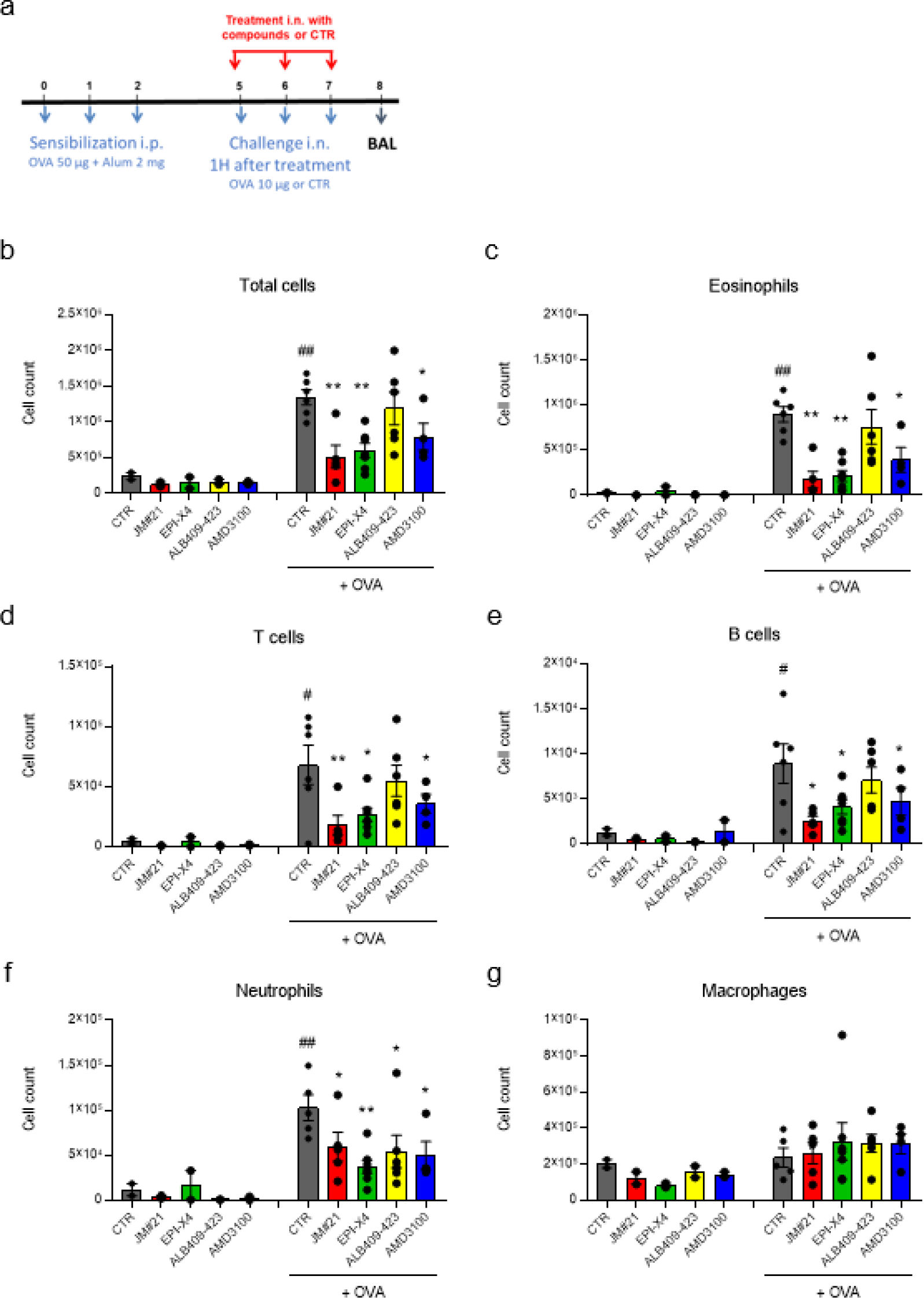
EPI-X4 JM#21 prevents immune cell infiltration into lungs in a mouse model of airway hypereosinophilia. (a) Balb/c mice were sensitized i.p. with a mixture containing 50 μg OVA and 2 mg alum in 0.2 ml saline on D0, D1 and D2 Mice were treated by i.n. administration with JM#21 (16 μmol/kg), EPI-X4 (16 μmol/kg), ALB409-423 (16 μmol/kg) or AMD3100 (12.6 μmol/kg), 2 hours before each OVA or saline challenge on D5, D6 and D7. N = 4-6 mice were used for OVA-challenged groups and n = 2 for unchallenged controls. Number of cells infiltrated into BAL were measured by flow cytometry, (b) all cells; (c) eosinophils; (d) T cells; (e) B cells; (f) neutrophils and (g) macrophages. Blocks are means and bars are SEM values. *# P* ≤ 0.05, ##*p* ≤ 0.01 compared to unchallenged control group, and * *p* ≤ 0.05, ** *p* ≤ 0.01 compared to CTR + OVA group.

### EPI-X4 JM#21 prevents skin inflammation in a murine model of atopic dermatitis

To evaluate the therapeutic potential of EPI-X4 JM#21 in an inflammatory skin disorder, we tested the peptide in a mouse model of atopic dermatitis. For this, ears of Balb/c mice were treated every 2 days with the vitamin D analog MC903 that induces changes in skin morphology and inflammation resembling immune perturbations observed in acute lesions of patients with AD^55^. EPI-X4 JM#21 and reference compounds AMD3100 and dexamethasone were applied topically daily (Fig. 9a). Thickness of ears was measured, and severity of the disease scored daily as described in Table 1. None of the compounds induced skin inflammation when administered in the absence of MC903 (Fig. 9b-d). At D11, MC903 induced a progressive and significant 3.1 ± 0.1-fold increase of the ear thickness (Fig. 9b,c), and a high disease score of 9.4 ± 0.1 (Fig. 9d) as compared to control (*p* < 0.0001). Treatment of the ears with JM#21 significantly inhibited ear thickening by 88 ± 4% (*p* < 0.0001), and the disease score by 88 ± 4 % (*p* < 0.0001). Thus, it was almost as effective as dexamethasone (96 ± 2 % and 97 ± 2, respectively, *p* < 0.0001) (Fig. 9b-d). The CXCR4 antagonist AMD3100 had only a minor effect on MC903 induced ear thickness (25 ± 7 % inhibition, *p* < 0.001) and disease score (18 ± 6 % inhibition, *p* < 0.001) (Fig. 9b-d). These results demonstrate that the optimized second-generation CXCR4 antagonist EPI-X4 JM#21 efficiently inhibits skin inflammation in a mouse model of dermatitis without causing any obvious side effects.

**Figure 9.**
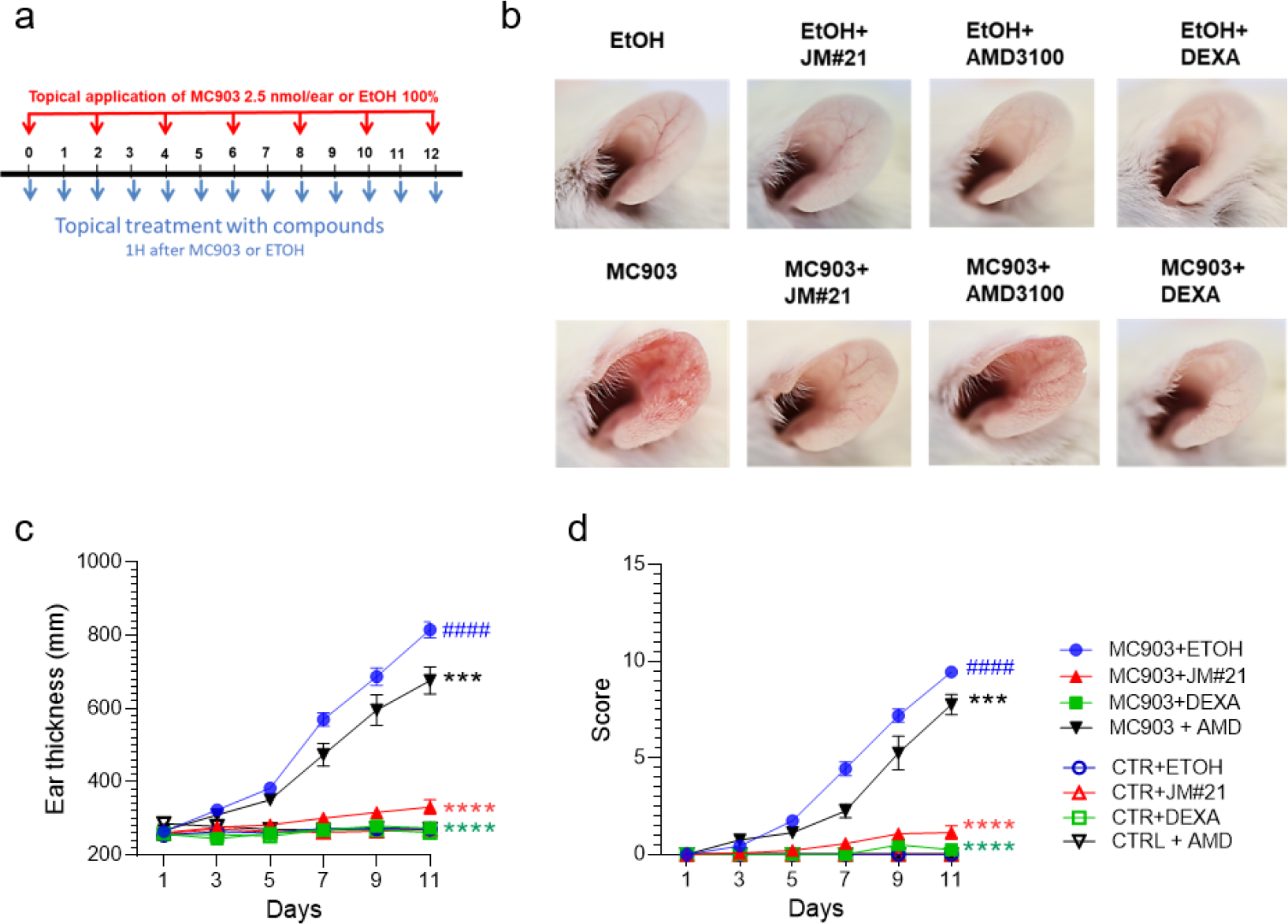
EPI-X4 JM#21 is active in a mouse model of atopic dermatitis. Ears of Balb/c mice were applied every two days with mock or MC903 (2.5 nmol/ear) (a). EPI-X4 JM#21 (400 nmol/ear), AMD3100 (315 nmol/ear), dexamethasone (Dexa) (20 nmol/ear) or ethanol (ETOH) were applied topically daily. Representative photographs of ears are taken at D12 (b). Ear thickness (c) and the inflammatory score (d) were determined daily. N = 8 - 16 mice were used for MC903 and n = 4 - 7 for controls (c + d). Data shown are mean values ± SEM. ####*p* ≤ 0.0001 compared to CTR + ETOH group, and \*\**p* ≤ 0.01, **** *p* ≤ 0.0001 compared to MC903 + ETOH group.

## Discussion

In this study we show, that rational drug design allowed to develop an optimized derivative of an endogenous CXCR4 antagonist that effectively prevents inflammatory reactions in murine models of atopic dermatitis and asthma. EPI-X4 JM#21 differs from WSC02, the first-generation derivative^34^, by three amino acid substitutions. Two exchanges were introduced because molecular docking studies suggested enhanced contacts of the peptide with the extracellular pocket in CXCR4. A third exchange was thought to achieve extra-flexibility of the mid part of JM21. Interestingly, NMR analyses revealed structural differences between JM#21 and WSC02, most notably the free N-terminal end in JM#21. This could explain the superior activity of JM#21 over WSC02 in cell-based assays, because binding with CXCR4 occurs in a random coil state of the peptide and as the peptide enters the CXCR4 core with its N-terminus (see Fig.1), where a free and accessible N-terminal end could be of advantage. Thus, a combination of empiric and rational drug design allowed to increase the anti-CXCR4 activity of EPI-X4 by nearly three orders of magnitude, as measured in the X4-HIV inhibition assay.

EPI-X4 JM#21 represents the lead of the 2^nd^ generation EPI-X4 derivatives. It inhibited CXCR4 antibody binding and X4-HIV infection, as well CXCL12-mediated ERK and AKT activation, Ca^2+^ mobilization and cell migration more efficiently than WSC02. JM#21 was also more active than clinically approved AMD3100, as determined in antibody competition, antiviral and ERK/AKT signaling assays. However, JM#21 was less active than AMD3100 in cell migration and Ca^2+^ mobilization assays. The reason(s) for these different activities in antagonizing CXCR4 remain to be determined and may reflect different binding modes of the small molecule AMD3100 versus the JM#21 peptide to CXCR4. Notably, none of the EPI-X4 derivatives induced acute, cardio- or developmental toxicity in zebrafish embryos, a widely accepted vertebrate model for toxicity assessment. Similarly, we did not observe visible or behavioral adverse effects of EPI-X4, WSC02 or JM#21 when administered topically onto the ears or intranasally into lungs of mice.

Intranasal application of JM#21 and EPI-X4 efficiently reduced influx of various immune and inflammatory cells but not macrophages into BAL in a mouse asthma model. Both peptides were more potent that AMD3100, which has previously been shown to attenuate allergen-induced infiltration of inflammatory cells into the airways^26,56,57^. The mechanism(s) underlying CXCR4-mediated inhibition of asthma are not fully understood, but may involve CXCR4-mediated suppression of inflammatory cell (mainly eosinophil) migration^22,24,25^, reduced expression of matrix metalloproteinase MMP-9 that drives airway remodeling^56^, or specific inhibition of CXCR4-expressing neutrophils or eosinophils releasing extracellular traps involved in asthma pathology^22,58^. Eosinophils are main contributors to the development of bronchial asthma and its exacerbations^54,58^. EPI-X4 and JM#21 reduced the number of eosinophils in lungs by 76 ± 7 % and 80 ± 1 %, respectively, suggesting a preferential application of the peptides in eosinophilic asthma. On average, application of 16 μmol/kg EPI-X4 and the optimized #JM21 analog suppressed inflammatory cell infiltration to a similar extent, which is not reflecting the different activities in antagonizing CXCR4. Perhaps both peptides have been applied at concentrations reaching maximum effects, suggesting that lower doses or less frequent application of JM#21 may result in similar outcomes, which will be analyzed in advanced preclinical development.

Notably, topical application of EPI-X4 JM#21 onto the ears effectively reduced skin inflammation in an atopic dermatitis mouse model. To our best knowledge, this is the first report demonstrating that a CXCR4-antagonizing agent shows therapeutic efficacy in an *in vivo* model of this inflammatory skin disease. Daily treatment with JM#21 reduced ear thickness and the disease score significantly more efficiently than AMD3100, and almost as efficiently as dexamethasone, an approved corticosteroid. The reason(s) for the differential efficacy of JM#21 and AMD3100 remain to be addressed. Differences in CXCR4 receptor binding and antagonization, or impaired penetration of AMD3100 into the skin, may be an explanation. It will be interesting to see whether a further increase of the JM#21 dose will result in a complete suppression of disease, as shown for dexamethasone. Thus, these findings warrant further development of JM#21 as therapeutic agent for atopic dermatitis, especially because the actual continuous treatment of inflamed skin with standard corticosteroids used nowadays is associated with side-effects and novel treatment options are urgently required^33^.

Our findings demonstrate that EPI-X4 and improved derivatives impair inflammatory reactions in mouse models of atopic dermatitis and asthma more efficiently than the FDA-approved small molecule AMD3100. It will be interesting to see, how EPI-X4 derivatives will perform in comparison to other classes of CXCR4 inhibitors, since several different molecules are currently under pre-clinical development for several different diseases with CXCR4/CXCL12 axis involvement. For JM#21, topical treatment of inflamed skin, or intranasal administration in the airway, might be an optimal application strategy, since it circumvents problems associated with peptide drugs, e.g. enzymatic instability and rapid excretion. Since, the CXCR4/CXCL12 axis is also involved in other inflammatory diseases of the skin and lungs (e.g. psoriasis^27^, and pulmonary fibrosis^59,60^), JM#21 might be an optimal drug candidate. In contrast to AMD3100, that is known for severe side effects after continues treatment and is therefore not approved for chronic diseases^14^, a peptide drug based on the body-own EPI-X4 will also most likely be well tolerated. Thus, further preclinical and clinical development of JM#21 as a novel class of CXCR4 inhibitors is highly warranted.

## Acknowledgements

This work was supported by the Deutsche Forschungsgemeinschaft (DFG) through the CRC1279 to L.S., G.K., E.S.-G., M.W., G.W., F.K. and J.M, the DFG by individual grants (MU 3115/11-1 and MU 3115(8-1) to J.M., the Baden-Württemberg Stiftung to J.M., and an ERC-PoC grant “EPI-X4 Health” to F.K and J.M. E.S.-G. was also supported by the DFG under Germany’s Excellence Strategy – EXC 2033 – 390677874 – RESOLV and by the Boehringer Ingelheim Foundation (Plus-3 Program).

